# The biochemical nature of senile plaques: intracellular destabilized-lysosome-enriched neural processes catabolizing extracellular hemorrhagic amyloid materials

**DOI:** 10.64898/2025.12.30.696965

**Authors:** Hualin Fu, Jilong Li, Chunlei Zhang, Guo Gao, Qiqi Ge, Daxiang Cui

**Affiliations:** School of Automation and Intelligent Sensing, Shanghai Jiao Tong University, Shanghai, China; Precision Research Center for Refractory Diseases, Institute for Clinical Research, Shanghai General Hospital, Shanghai Jiao Tong University School of Medicine, Shanghai, China

**Keywords:** Alzheimer’s disease, senile plaque, Aβ, Tau phosphorylation, lysosome destabilization, blue autofluorescence, MAP2, hemorrhage, microaneurysm, Cathepsin D

## Abstract

The pathological mechanism of senile plaque formation in Alzheimer’s disease remains a challenging unresolved problem for over one hundred years. In this study, we investigated senile plaque pathogenesis based on immunohistochemistry, fluorescence imaging and quantitative analysis of AD front brain sections with multiple senile plaque related markers such as Aβ, MAP2, phos-Tau, Cathepsin D, Sortilin 1, Ceruloplasmin and Hemin. Firstly, we confirmed that senile plaque Aβ deposition mostly happened in MAP2-positive but degenerative neuritic processes. Secondly, we found an interesting dual-component granular or diffusive staining pattern of all of these senile plaque related markers, matching to two states of lysosomes, relatively-stable granule lysosomes and destabilized diffusive lysosomes as shown by lysosome-related marker Cathepsin D or Sortilin 1 staining. Thirdly, dystrophic senile plaque neural processes contain hemorrhage-related markers such as Ceruloplasmin and Hemin. In addition, Aβ deposition, Tau phosphorylation, lysosome destabilization are all enriched at the sites of microaneurysm formation. Moreover, senile plaques are also characterized by a widespread binary phos-Tau staining pattern with difference in term of staining intensity. We concluded that senile plaques are formed by destabilized-lysosome-enriched neural processes catabolizing extracellular hemorrhagic amyloid materials, with microaneurysm as an important intermediate stage.

## Introduction

The pathological hallmarks of Alzheimer’s disease include senile plaques, neurofibrillary tangles and neuritic dystrophy[1, 2]. In spite of countless research on Alzheimer’s disease for more than one century, the pathological mechanism of senile plaque formation in AD is not completely understood, highlighting the tremendous complexity of this disease[3–5]. There is intensive debate on whether senile plaque amyloid materials come from neural cells[6, 7], glial cells[8, 9], vascular cells or the blood[10–14]. A dozen of interconnecting, non-exclusive models have been proposed to explain the cause of neurodegeneration in AD such as the amyloid cascade model, which suggests that Aβ deposition causes neurofibrillary tangle formation and neuronal cell death[6], synaptic toxicity theory [15, 16], mitochondria deficiency theory [17, 18], autophagy and lysosome defects[19–22], neural inflammation [23, 24], metal ion toxicity such as iron, copper or aluminum toxicity [25–27], oxidative stress [28, 29], DNA damage[30, 31], vascular defect [32, 33], cholesterol overload[34, 35] etc., while a clear integrated mechanism of AD pathogenesis is still lacking. Although the AD research and therapy field has advanced so much to come up several medicines for AD disease modification therapy such as lecanemab[36] and donanemab[37], even without a complete understanding of AD pathology, an integrated pathological mechanism for AD is urgently needed in the fight against AD pandemic.

Multiple studies indicate that lysosome proteins are enriched in or around senile plaques during Alzheimer’s disease pathogenesis[20, 38–40]. In addition, massive accumulation of luminal protease-deficient axonal lysosomes at amyloid plaques was identified[21]. However, how lysosome defects relate to Aβ deposition, tau phosphorylation and senile plaque formation is not completely understood.

Over the past several years, our group established a linkage between senile plaque formation and microaneurysm rupture[14]. We provided solid evidence that the cores of dense-core plaques don’t have nuclei staining but instead contain blood and vascular-derived materials such as Hemin, HbA1C, ColIV and wrapped with glial foot marker GFAP, suggesting the cores of dense-core plaques are of vascular and blood origin. We also characterized the mysterious blue autofluorescence in the senile plaques as an intrinsic property of Aβ self-oligomerization or hetro-oligomerization with Hemoglobin[41]. Further, our studies suggested that pathological axonal enlargement in connection with amyloidosis, lysosome destabilization and hemorrhage is a major defect in Alzheimer’s disease[42]. However, due to the limitation of the previous study which focused on the cores of dense core senile plaques[14], the complete process of senile plaque formation, especially the formation of diffusive plaques and the diffusive corona portion of dense-core plaques still remains obscure. In this study, we aimed to obtain a complete view of senile plaque pathogenesis by examining the mechanism of diffusive senile plaque development through immunohistochemistry and fluorescence imaging analysis, emphasizing the role of lysosome destabilization during diffusive senile plaque formation. The data came up a very simple model that senile plaques are mainly formed by destabilized-lysosome-enriched neural processes internalizing extracellular hemorrhagic amyloid materials.

## Material and methods

### Tissue sections

AD patient frontal lobe brain paraffin tissue sections were purchased from GeneTex (Irvine, CA, USA). Additionally, AD patient and non-AD control frontal lobe brain paraffin tissue sections were provided by National Human Brain Bank for Development and Function, Chinese Academy of Medical Sciences and Peking Union Medical College, Beijing, China. This study was supported by the Institute of Basic Medical Sciences, Chinese Academy of Medical Sciences, Neuroscience Center, and the China Human Brain Banking Consortium. All experiment procedures were performed in accord with the ethical standards of the Committee on Human Experimentation in Shanghai Jiao Tong University and the Helsinki Declaration of 1975.

### List of antibodies for immunohistochemistry

The following primary antibodies and dilutions have been used in this study: Aβ (Abcam ab201061, 1:200, Cambridge, UK), Aβ/AβPP (CST #2450, 1:200, Danvers, MA, USA), phos-Tau (Abcam ab151559, 1:200), MAP2 (Proteintech 17490-1-AP, 1:200, Rosemont, IL, USA), Sortilin1 (Abcam ab263864, 1:200), HBA (Abcam ab92492, 1:200), Cathepsin D (Abcam ab75852, 1:200), Lamp2 (Abcam ab199946, 1:200), Cathepsin D (Proteintech 66534-1-Ig, 1:100), Ceruloplasmin (Proteintech, 21131-1-AP, 1:200), Hemin (Absolute Antibody, 1D3, 1: 100, Redcar, UK), ApoE (Abcam ab183597, 1: 200), HbA1c (OkayBio K5a2, 1: 100, Nanjing, Jiangsu, China). The following secondary antibodies and dilutions have been used in this study: donkey anti-mouse Alexa-594 secondary antibody (Jackson ImmunoResearch 715-585-150, 1:400, West Grove, PA, USA), donkey anti-rabbit Alexa-488 secondary antibody (Jackson ImmunoResearch 711-545-152, 1:400), donkey anti-rabbit Alexa-594 secondary antibody (Jackson ImmunoResearch 711-585-152, 1:400), and donkey anti-mouse Alexa-488 secondary antibody (Jackson ImmunoResearch 715-545-150, 1:400).

**Immunohistochemistry** was performed as described[43]. Briefly, paraffin sections were deparaffinized by Xylene, 100% EtOH, 95% EtOH, 75% EtOH, 50% EtOH, and PBS washes. Sections were then treated with 10 mmol/L pH 6.0 sodium citrate or 10 mmol/L pH 9.0 Tris-EDTA antigen retrieval solutions in a microwave at high power for 5 minutes to reach the boiling point and then at low power for another 15 minutes. The sections were allowed to naturally cool down to room temperature. Then, the slides were blocked with TBST 3% BSA solution for 1 hour at room temperature. After blocking, the samples were incubated with primary antibodies at room temperature for 2 hrs followed by 5 washes of TBST. After that, the samples were incubated with fluorescent secondary antibodies overnight at 4 degree. The treated samples were washed again with TBST 5 times the second day and mounted with PBS+50% glycerol supplemented with Hoechst nuclear dye (Sigma, B2261, 1 μg/ml) and ready for imaging. IHC experiments without primary antibodies were used as negative controls. All experiments have been repeated in order to verify the reproducibility of the results.

### Imaging and morphometry analysis

The fluorescent images were captured with a CQ1 confocal fluorescent microscope (Yokogawa, Ishikawa, Japan). Images were then analyzed with ImageJ software (NIH, USA). When comparing signal intensities with the same exposure settings, the mean area density was used as the parameter to define the marker densities. When measuring the intensities of regions of interests comparing to their backgrounds, freehand selection method was used to define selected ROIs. “Make reverse” command under the “Selection” submenu was used to select the “background” region of the defined ROI. The average area intensities of different regions were then compared.

### Colocalization analysis

Quantitation of the colocalization of paired markers was performed with the Image J “Colocalization Threshold” plugin. First, the interested area was selected with a small rectangle. To increase the sensitivity of the analysis, the interested object could be further defined with a freehand selection ROI (region of interest). Paired RGB images from two different fluorescent channels were converted into 16-bit images and then analyzed with the “Colocalization Threshold” plugin with the ROI selection. The quantitation of colocalization was reported as M1 after threshold (tM1) or tM2 for the two respective markers. A merged picture highlighting the colocalized pixels was also produced by the software.

### Statistics

All data first went through a Shapiro & Wilk normality test using SPSS Statistics 19 software. Two-tailed unpaired T-test was used to compare the means of data with a normal distribution with Excel 2007 software. For data that were not normally distributed, nonparametric Mann-Whitney test was performed to compare the means by using SPSS Statistics 19 software. The p Value threshold for statistical significance is set at 0.05. If p<0.05, then the difference was considered statistically significant. When p<0.001, it was labeled as p<0.001. If p≧0.001, then the p Value was shown as it was.

## Funding

This work was supported by the National Natural Science Foundation of China (81472235), the Shanghai Jiao Tong University Medical and Engineering Project (YG2021QN53, YG2017MS71), the International Cooperation Project of National Natural Science Foundation of China (82020108017), and the Innovation Group Project of National Natural Science Foundation of China (81921002), National Key Research and Development Program of China (No. 2017FYA0205301).

## Author contributions

HF conceived the study, designed and supervised the experiments and wrote the manuscript; HF, JL performed the experiments; HF, JL, CZ, GG, QG and DC did the data analysis; all authors reviewed the manuscript.

## Competing interests

Authors declare no competing interests.

## Data availability statement

The data that supports the findings of this study are available from the main manuscript and the supplementary files.

## Supplementary Materials

Supplementary Figures 1 to 7

## Results

### Senile plaques are mainly consisted of Aβ-positive degenerative neuronal processes with drastically reduced MAP2 staining

Aβ is known as one of the most important molecules involving in Alzheimer’s disease pathogenesis. How Aβ triggers neurodegeneration in AD is not completely clear. In our previous studies[42], the data showed that Aβ deposition significantly reduces MAP2 expression in neuronal processes in AD brains while senile plaque regions appear as “black holes” with a lack of MAP2 expression. It is not clear how neural processes degenerate within senile plaques due to the difficulty of imaging the weak residual MAP2 signals. In this study, we further optimized MAP2 detection parameters in the senile plaques in order to image the detail phenotypes of amyloidosis-related neuritic MAP2 degeneration. We found that neuronal processes in senile plaque regions in AD brains have drastically reduced MAP2 expression with a distinct beading phenotype (Figure 1). Senile plaque Aβ deposition actually runs along the neuronal processes with the beading type of MAP2 staining. MAP2 staining in senile plaques was significantly reduced compare to proximal neighboring brain regions outside of senile plaques. The data showed that senile plaque region has significantly reduced MAP2 intensity, which is around 0.69±0.08 (N=8) of the average MAP2 intensity level of the immediate surrounding region in AD brain tissues. In another word, neuronal processes in the senile plaque region lost roughly 31% of MAP2 expression comparing to the surrounding AD brain tissues. As we know from previous studies, MAP2-positive axon number in axon bundles is reduced by more than half in AD brains comparing to control brains[42]. The new data suggested that the senile plaque regions have further reduction of MAP2 levels even comparing to neighboring regions in AD brain tissues. The immediate effect of Aß deposition and senile plaque formation on neurodegeneration appears to be the neuronal process degeneration within senile plaques, which can be demonstrated clearly by MAP2 reduction.

**Figure 1.**
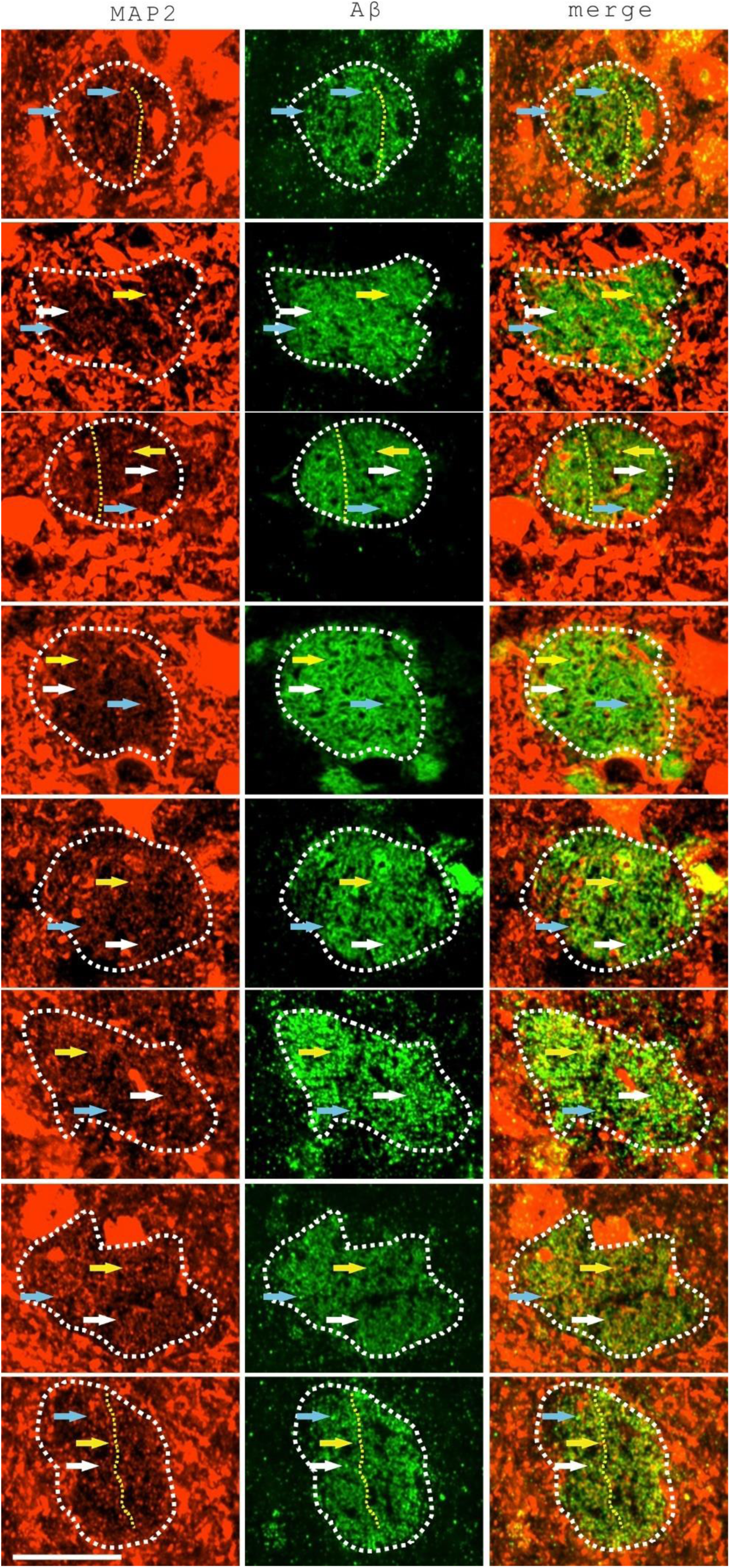
Senile plaque development colocalized with degenerative neuronal processes with weak MAP2 staining and a severe beading phenotype. Eight representative images were shown. Quantitation of the intensity of MAP2 in senile plaques vs. regions outside of senile plaques showed that senile plaque region has greatly reduced MAP2 intensity, which is around 0.69±0.08 (N=8) comparing to the immediate surrounding region MAP2 levels. The yellow arrows indicated the beading-type MAP2 staining of the degenerative neuronal processes. The white arrows indicated the weak diffusive type of MAP2 staining in degenerating neurites. The white dashed circles indicated senile plaques. The yellow dashed lines indicate several relatively long neurites with both Aβ deposition and beading-type degenerative MAP2 staining. The blue arrows indicated granule-type Aβ staining in the senile plaques. Scale bar, 25 µm.

### Senile plaques can be visualized by endo-lysosomal markers such as Cathepsin D and Sortilin1, with signs of lysosome destabilization

How amyloid deposition in senile plaques causes MAP2 reduction? We previously proposed that lysosome destabilization could mediate the effect of Aβ on MAP2 reduction in axons and neuronal soma[42]. In this study, we further optimized the parameters of lysosome detection in senile plaques with two different lysosome-related markers, Cathepsin D (Figure 2) and Sortilin1 (Figure 3). The data showed that senile plaques could actually be imaged with lysosomal markers with high resolution since Aβ staining in senile plaques highly co-localized with lysosome markers such as Cathepsin D and Sortilin 1. Elevated Cathepsin D staining (1.80±0.30, N=9, p<0.001, T-test) in senile plaques was detected comparing to the immediate surrounding region. Sortilin1 staining in senile plaques was also enhanced comparing to neighboring regions, which is around 1.75±0.29 (N=9, p<0.001, T-test) times over immediate background Sortilin 1 staining. Colocalization quantitation showed that approximately 69±21% (N=9) of Aβ staining colocalized with Cathepsin D staining in senile plaques while approximately 67±17% (N=9) of Aβ staining colocalized with Sortilin1 staining in senile plaques. Examples of colocalization analysis between Aβ and Cathepsin D or Sortilin 1 by ImageJ Colocalization Threshold plugin were provided in Supplementary Figure 1. However, both Cathepsin D and Sortilin1 labels SP with less contrast comparing to Aβ. The enhancement of Aβ in senile plaques reached 7.21±2.40 (N=18) times comparing to average Aβ staining intensity of the immediate neighboring regions.

**Figure 2.**
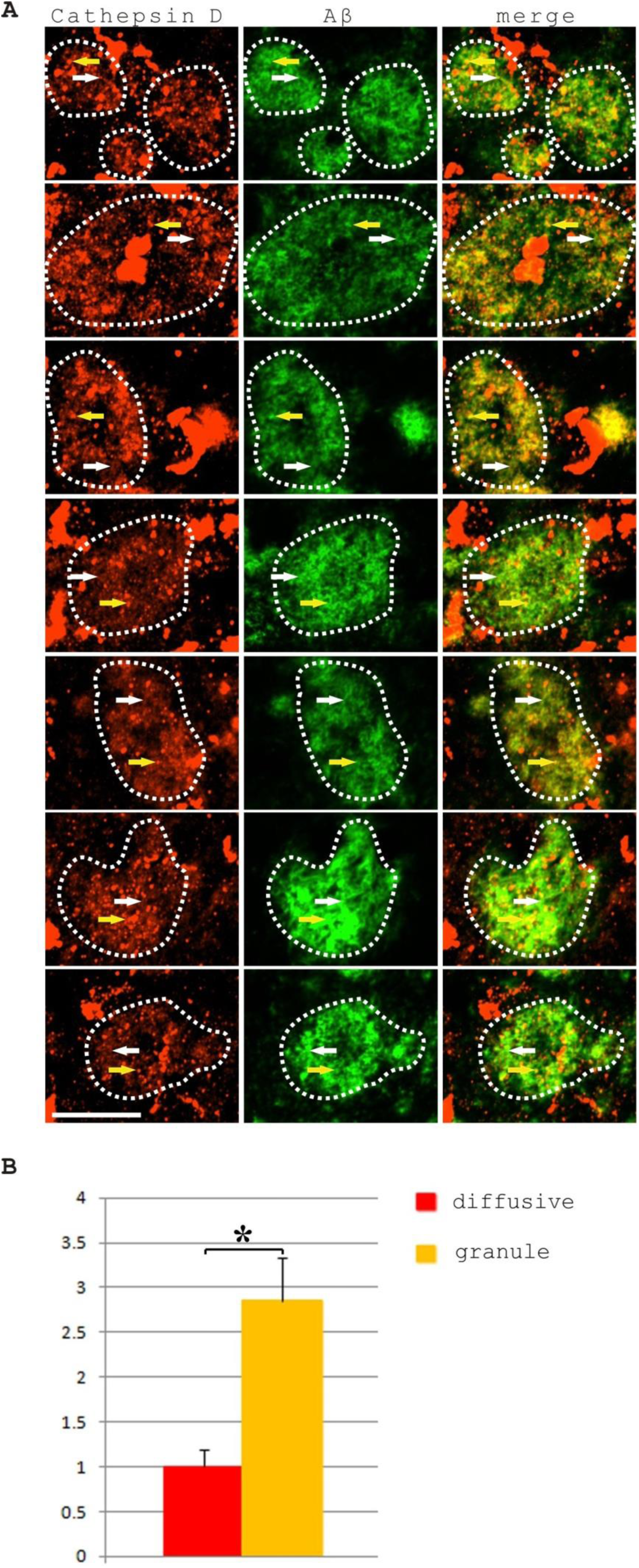
Senile plaques can be labeled by Cathepsin D staining clearly. A, Senile plaque Aβ immunostaining colocalized with granule and diffusive staining patterns of lysosome marker Cathepsin D. Seven representative images were shown. The yellow arrows indicated the granule-type Cathepsin D. The white arrows indicated the weak diffusive type of Cathepsin D staining in degenerating neurites. The white dashed circles indicated senile plaques. Measurements showed that approximately 29±8% (N=9 senile plaques) of the Cathepsin D staining was in the granule-stained lysosomes while the vast majority of staining 71% was in the diffusive pattern. Comparatively, 15±4% (N=9 senile plaques) of the Aβ staining was in the granule-stained lysosomes while the majority Aβ of staining around 85% was in the compartments with diffusive Cathepsin D staining. Colocalization quantitation showed that approximately 69±21% (N=9) of Aβ staining colocalized with Cathepsin D staining in senile plaques. Scale bar, 25 µm. B, The average Cathepsin D staining intensity of granule-like lysosomes is 2.85±0.49 (N=466 lysosomes, p<0.001, Mann-Whitney test) times of Cathepsin D staining intensity in the diffusive destabilized lysosomal compartments. Aβ staining intensities in the Cathepsin D-labeled granule-like lysosomes are also significantly stronger (1.36±0.58, N=466, p=0.049, Mann-Whitney test) than Aβ staining intensities in the regions with diffusive Cathepsin D staining.

**Figure 3.**
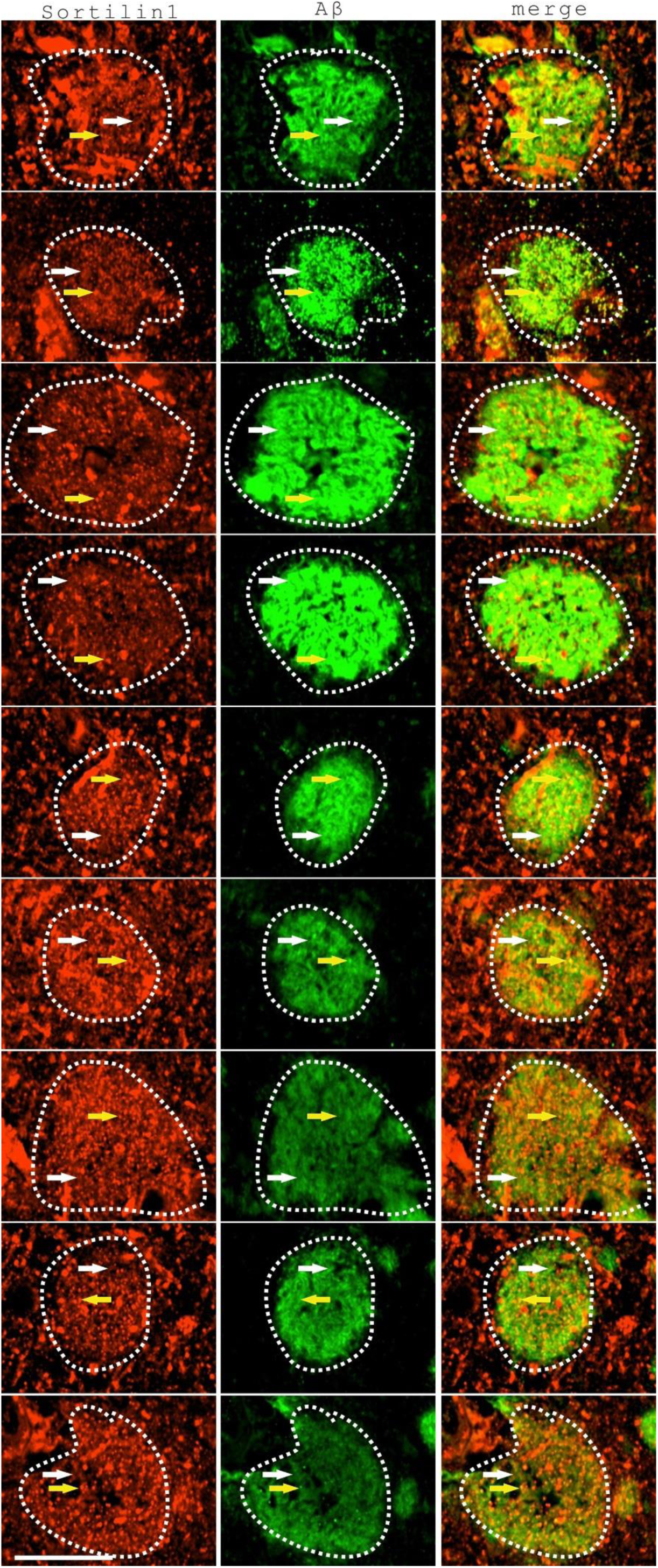
Senile plaques colocalized with granule and diffusive staining patterns of endo-lysosome marker Sortilin1. Nine representative images were shown. The yellow arrows indicated the granule-type Sortilin 1 staining. The white arrows indicated the weak diffusive type of Sortilin 1 staining in degenerating neurites. The white dashed circles indicated senile plaques. Measurements showed that approximately 31±12% (N=9) of the Sortilin1 staining was in the granule-stained endo-lysosomes while the vast majority of staining roughly 69% was in the diffusive pattern. Comparatively, 21±9 % (N=9) of the Aβ staining was in the granule Sortilin1-stained endo-lysosomes while the majority of Aβ staining (approximately 79%) was in the compartments with diffusive Sortilin1 staining. Colocalization quantitation showed that approximately 67±17% (N=9) of Aβ staining colocalized with Sortilin1 staining in senile plaques. Scale bar 25 µm.

After examining staining patterns of Cathepsin D and Sortilin1 in senile plaques carefully, we noticed that, just like Aβ staining pattern in Figure 1, both Cathepsin D and Sortilin 1 staining showed a characteristic dual-component expression pattern in the senile plaques, the granule staining pattern of typical intracellular vesicles and the diffusive staining pattern along the neural fibers. Our previous study suggested that Aβ along with other hemorrhagic amyloid materials in the lysosomes might induce lysosome destabilization with lysosome clustering, enlargement and possibly permeabilization, which could be related to the diffusive appearance of lysosome markers[42]. The diffusive staining along the neuronal processes in senile plaques very likely reflects lysosome destabilization events associating with Aβ lysosomal toxicity. Measurements showed that, in senile plaques, approximately 29±8% (N=9) of the Cathepsin D staining (measured by area intensity) was in the granule-stained lysosomes while the vast majority of Cathepsin D staining (around 71%) was in the diffusive pattern. Comparatively, 15±4% (N=9) of the Aβ staining was in the granule-stained Cathepsin D-labeled lysosomes while the majority of Aβ staining (85%) was in the compartments with diffusive Cathepsin D staining.

The analysis on the Sortilin 1 staining yielded similar results. Measurements showed that approximately 31±12% (N=9) of the Sortilin1 staining was in the granule-type Sortilin 1-stained endo-lysosomes while the vast majority of staining roughly 69% was in the diffusive pattern. Comparatively, 21±9% (N=9) of the Aβ staining was in the granule-type Sortilin 1-stained endo-lysosomes while the majority of Aβ staining (approximately 79%) was in the compartments with diffusive Sortilin1 staining. In addition, there was a clear intensity difference of Cathepsin D or Sortilin 1 in granule-like lysosomes vs. diffusive lysosome staining, which demonstrate an overall morphology resembling “sesame cookies”, with granule particles resembling sesame seeds while the diffusive approximately round-shape senile plaque staining forms the base of “cookies”. The average Cathepsin D staining intensity of granule-like lysosomes is 2.85±0.49 (N=466 lysosomes, p<0.001, Mann-Whitney test) times of Cathepsin D staining intensity in the diffusive destabilized lysosomal compartments. The average Sortilin 1 staining intensity of granule-like vesicles is 1.81±0.27 (N=833 granules, p<0.001, Mann-Whitney test) times of Sortilin 1 staining intensity in the diffusive destabilized endo-lysosomal compartments. However, Aβ staining intensity in the Cathepsin D-labeled granule-like lysosomes showed less but still significant enhancement (1.36±0.58, N=466, p=0.049, Mann-Whitney test) over Aβ intensity in the regions with diffusive Cathepsin D staining in senile plaques. This means Aβ staining intensity is relatively higher in granule-type lysosomes than diffusively-stained lysosomes. However, overall, granule-type Aβ staining pattern is less-frequently observed comparing to Cathepsin D or Sortilin 1 granule staining in senile plaques.

Results above indicated that the majority of Aβ existed in destabilized lysosomal compartments colocalizing with diffusive lysosome staining marked with lysosome-related markers such as Cathepsin D or Sortilin 1. Lysosome destabilization with leaky diffusive protease degradation could be directly related to the MAP2 degradative phenotype in the senile plaque neurites as also suggested in the previous study [42], implicating proteomic damage leading to neuronal process degeneration is a closely-linked direct downstream event of Aβ deposition in AD.

### Senile plaques bear the markers of hemorrhage such as Ceruloplasmin and Hemin

Does the endogenous intraneuronal Aβ cause the senile plaque formation or senile plaques are formed by neurites intaking extracellular Aβ and other associated toxic materials from hemorrhagic leakage? The evidence from our previous studies [14, 41] suggested that Aβ in the cores of dense-core senile plaques was mostly derived from an extracellular source of hemorrhagic leakage since Aβ in the dense-core is associating with many hemorrhagic markers such as Hemin and HbA1C. In this study, two markers of blood-related materials, Ceruloplasmin and Hemin, demonstrated that the neuronal processes in the diffusive senile plaques were also enriched for hemorrhagic materials (Figure 4). Both Ceruloplasmin and Hemin showed similar “sesame cookie” type of staining patterns as Cathepsin D and Sortilin1, with clear intensity difference between the granule pattern vs. the diffusive type pattern. The intensity of Ceruloplasmin staining in the granule pattern is 2.83 (2.83±0.48, N=745 granules, p<0.001, Mann-Whitney test) times of the Ceruloplasmin staining intensity in the diffusive pattern. The intensity of Hemin staining in the granule pattern is 2.18 (2.18±0.30, N=543 granules, p<0.001 Mann-Whitney test) times of the Hemin staining intensity in the diffusively-stained regions within senile plaques. Colocalization quantitation showed that 89±7% (N=9 plaques) of Ceruloplasmin colocalized with Aβ staining in senile plaques while 72±14% (N=6 plaques) of Hemin staining colocalized with Aβ staining in senile plaques. These results together with previous study [14] indicated that Aβ is strongly co-localized with hemorrhagic markers in both dense-core senile plaques and diffusive senile plaques.

**Figure 4.**
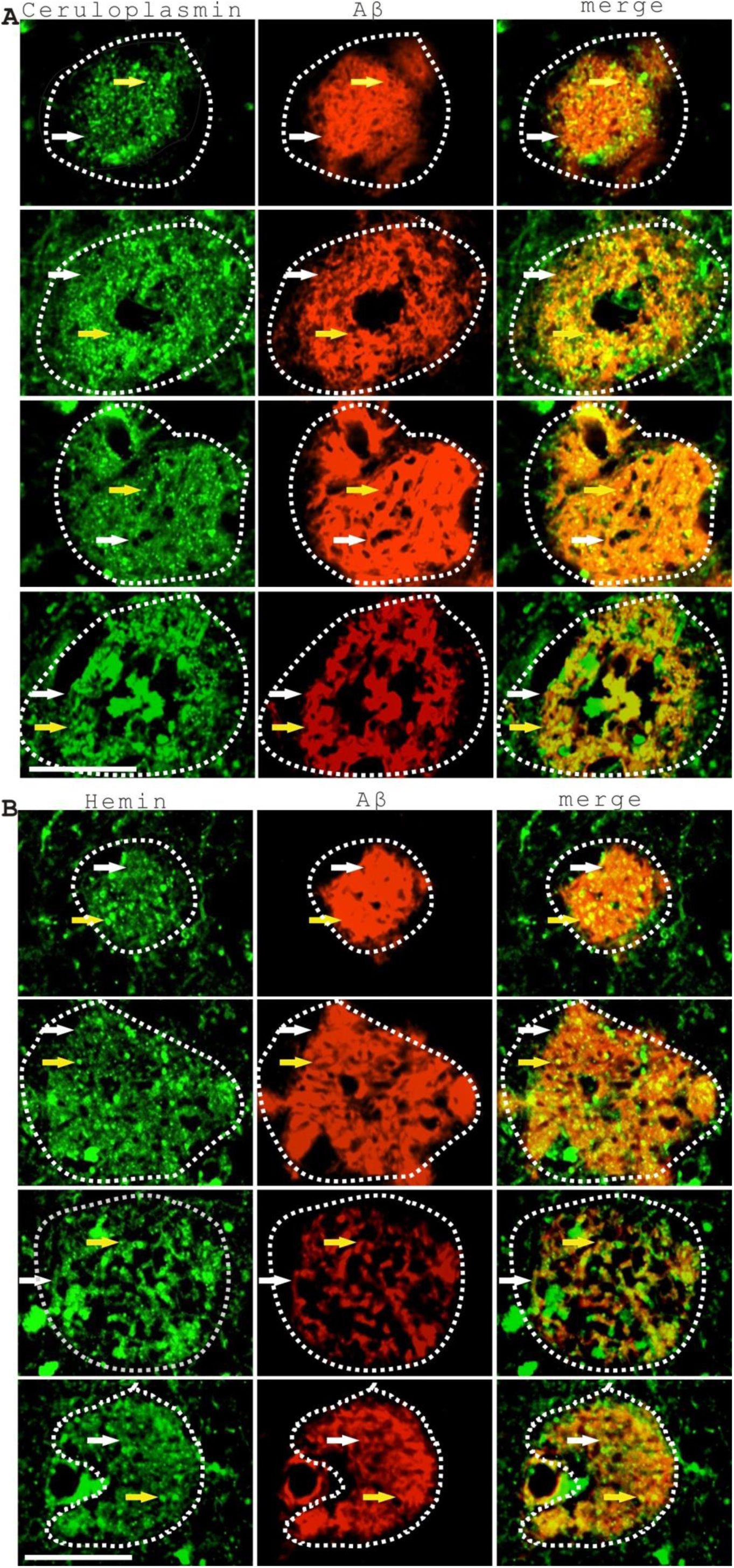
Senile plaque Aβ staining colocalized with granule and diffusive staining patterns of hemorrhagic marker Ceruloplasmin and Hemin. (A) senile plaque Aβ (red) colocalized with granule and diffusive staining patterns of hemorrhagic marker Ceruloplasmin (green). The yellow arrows indicated the granule-type Ceruloplasmin staining. The white arrows indicated the weak diffusive type of Ceruloplasmin staining in degenerating neurites. The white dashed circles indicated senile plaques. Colocalization quantitation showed that 89±7% (N=9 plaques) of Ceruloplasmin colocalized with Aβ staining. Scale bar 25 µm. (B) Senile plaque Aβ (in red) colocalized with granule and diffusive staining patterns of hemorrhagic marker Hemin (in green). The yellow arrows indicated the granule-type Hemin staining. The white arrows indicated the weak diffusive type of Hemin staining in degenerating neural processes. The white dashed circles indicated senile plaques. Colocalization quantitation showed that 72±14% (N=6 plaques) of Hemin staining colocalized with Aβ staining. Scale bar 25 µm.

### Microaneurysm rupture induces neurodegeneration and serves as an intermediate stage of senile plaque formation

If senile plaques derive from hemorrhage events, why we did not observe bleeding in the senile plaques more frequently? The reason could be that senile plaque formation goes through an intermediate stage of microaneurysm formation and rupture in a chronic, silent, hardly-noticable fashion. The frequency of microaneurysms is highly variable in end-stage brain tissues in AD patients ranging from several to hundreds on different pathological sections[14]. Careful examination of many microaneurysms suggested that neuronal process MAP2 marker degeneration already took place at the stage of microaneurysm rupture and stressed neuronal process already interacted with microaneurysms at the early stage of senile plaque pathogenesis (Figure 5). The neural processes interacting with microaneurysms could be either from the vascular-innervating neurons regulating blood vessels as suggested previously[4] or neural processes of other neurons in the brain parenchyma that have been pushed against the microaneurysms pathologically during the cerebral microvascular malformation. Beading-type degenerative MAP2 staining was observed in the neurites associating with microaneurysm. In the microaneurysms, Aβ was also clearly enriched but with a relatively diffusive pattern. The average Aβ staining intensity is around 3.03±0.97 (N=5, p=0.009, Mann Whitney test) times of the average Aβ intensity of the immediate sample environment. In addition, when we analyzed another marker of neurodegeneration, phosphorylated Tau, we could also detect the enrichment of Tau phosphorylation specifically surrounding microaneurysm ruptures (Figure 6). Tau phosphorylation is specifically enriched at around 2.45±0.57 (N=7, p<0.001, T-test) times of the average intensity of immediate surrounding tissue environment as shown in Figure 6. The increase of Tau phosphorylation in microaneurysms is also associated with the increase of lysosome destabilization shown by diffusive Lamp2 immunostaining. The overall intensity of Lamp2 staining was also enhanced (3.38±1.08, N=7, p<0.001, T-test) comparing to the immediate environment of microaneurysm rupture. However, Lamp2 staining surrounding microaneurysms were not normal looking, but instead, showed a blend of diffusive Lamp2 staining and granule type Lamp2 staining, very similarly as the lysosomal staining in senile plaques. This indicated that lysosomal destabilization was already happening at the microaneurysm stage. Although some residual Lamp2 staining in the microaneurysms might be inherited from degrading lysosomal materials during the blood or vascular inflammation as we previously suggested[14], abundant lysosome organelles in senile plaques are likely de-novo generated and recruited in responding to the hemorrhagic leakage insults by the neurons. The enrichment of amyloidosis-related materials in the microaneurysm could also be indicated with amyloid-related blue autofluorescence (Figure 6, the blue fluorescence panel). We noticed that microaneurysms were also enriched with some black materials (Figure 6, yellow arrows), which showed dark staining under phase contrast microscopy. In our previous studies, these black materials co-distributed with blood markers such as Hemin, which could be signs of hemorrhage. These results together suggested that amyloid material leakage from microaneurysm rupture induces lysosome destabilization, MAP2 degradation and Tau phosphorylation in the microaneurysm lesions, very similarly as what happened in the senile plaques. At the end stage of AD patients, probably most senile plaques have gone through the microaneurysm formation and rupture stage already, so that the microaneurysm stage of senile plaque formation in AD was only infrequently noticed.

**Figure 5.**
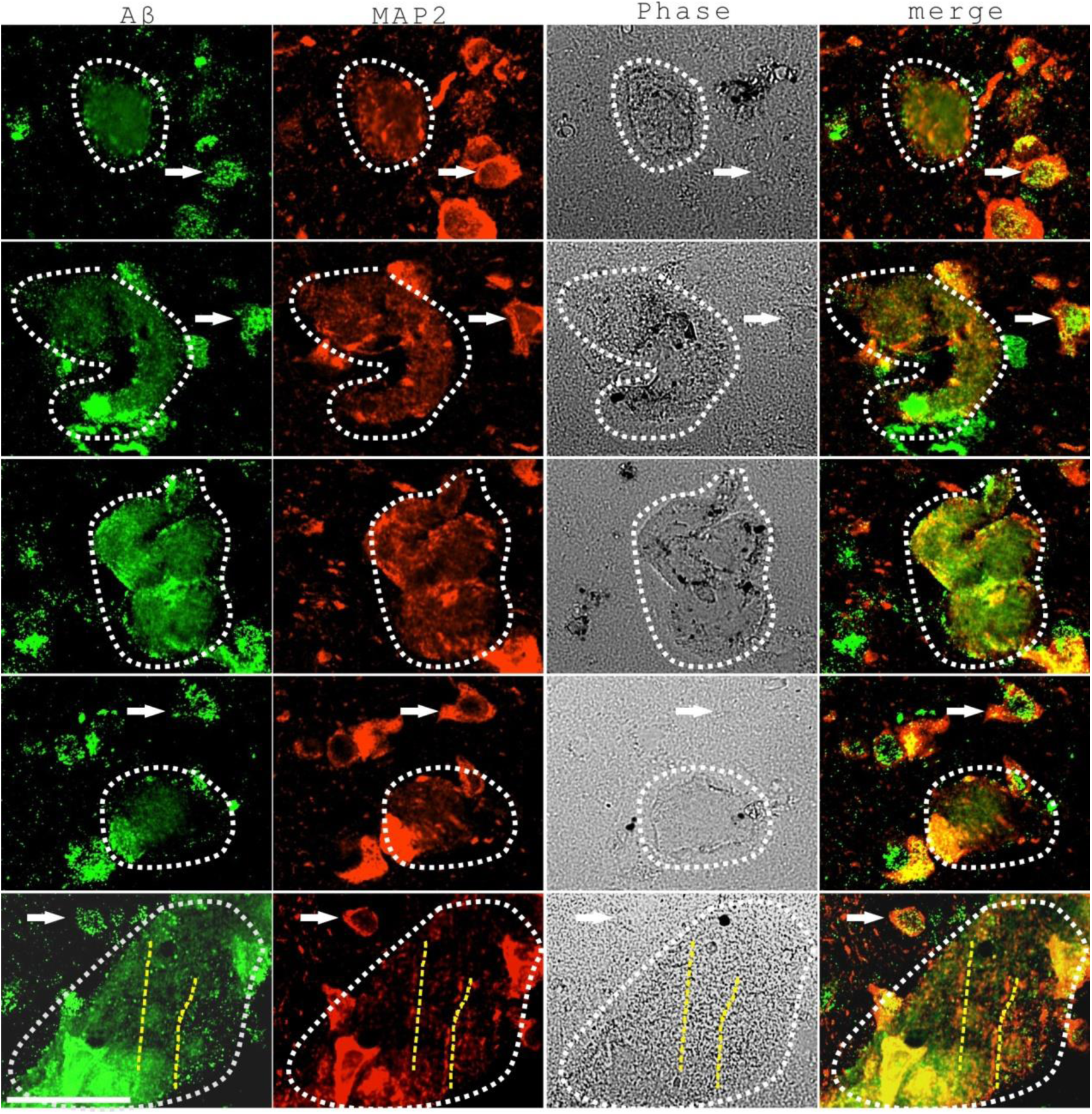
Microaneurysms induced Aβ deposition and MAP2 degeneration in neurites. Five samples were shown. Aβ was stained with green fluorescence while MAP2 was stained with red fluorescence. Microaneurysms were clearly identified with Phase Contrast microscopy. The average Aβ staining intensity in microaneurysms is around 3.03±0.97 (N=5) times of the average Aβ intensity of the immediate tissue environment. White dashed lines indicated the regions of microaneurysms. White arrows point to neighboring neurons with MAP2 immunostaining. Yellow dashed lines indicated two long neurites showing beading type degenerative MAP2 staining. Scale bar, 50 µm.

**Figure 6.**
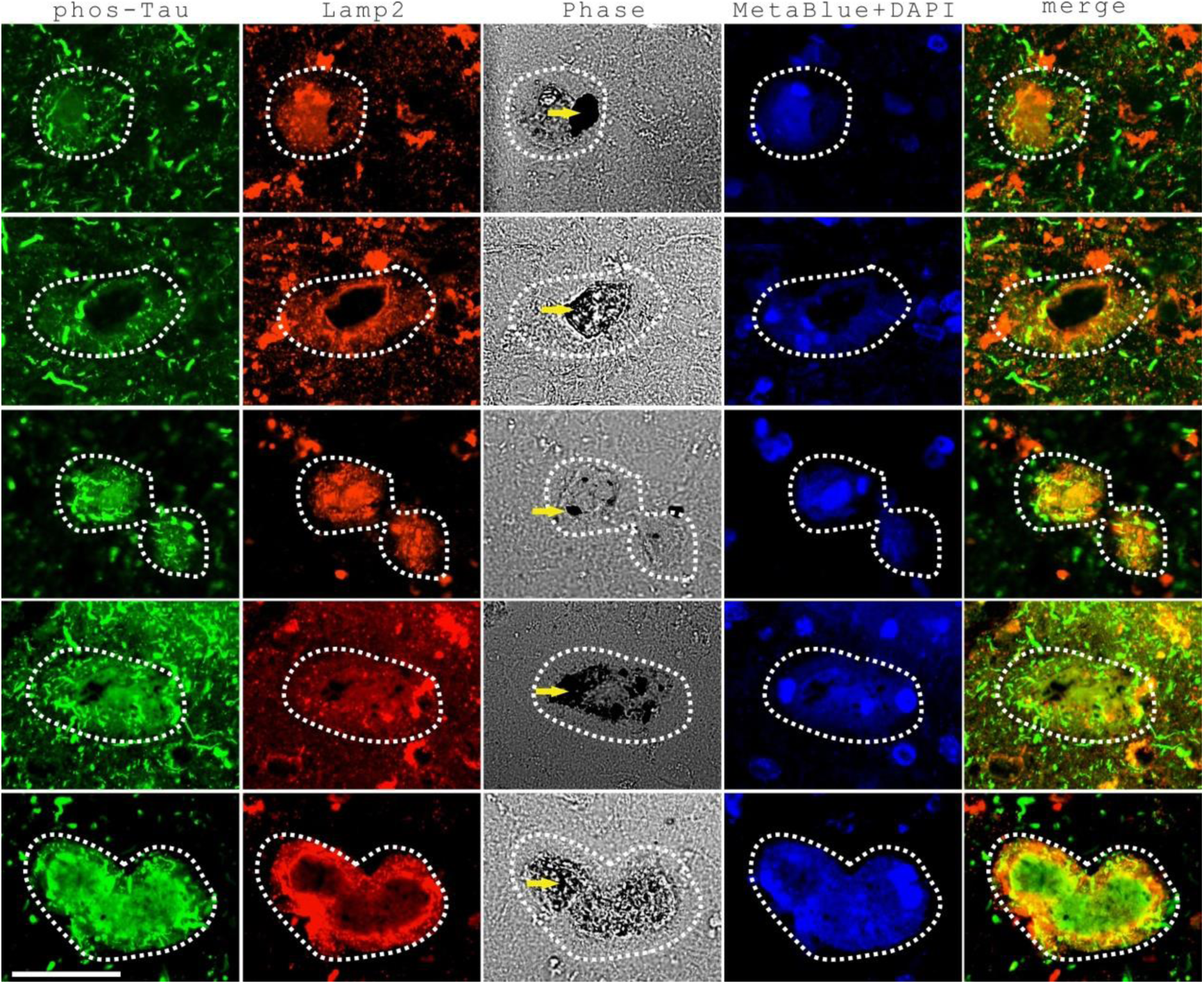
Microaneurysms were the converging sites of Tau phosphorylation, lysosome destabilization and amyloid deposition. Five representative microaneurysms were shown. Lysosome marker Lamp2 was stained with red fluorescence while phos-Tau was stained with green fluorescence. Microaneurysms were identified with Phase Contrast microscopy. Microaneurysm amyloidosis was also indicated with amyloid blue autofluorescence. Tau phosphorylation is specifically enriched in and around the microaneurysm sites with higher Tau phosphorylation intensity at around 2.45±0.57 (N=7) times of the average intensity of immediate tissue environment as shown in Figure 6. The increase of Tau phosphorylation surrounding microaneurysms is also associated with the increase of lysosome destabilization shown by diffusive Lamp2 immunostaining. The overall intensity of Lamp2 staining was also enriched comparing to the immediate environment of microaneurysm rupture (Lamp2 staining intensity around microaneurysms is around (3.38±1.08, N=7) times of the average intensity of the immediate background. White dashed lines indicated the microaneurysms. Yellow arrows indicated some black materials in the microaneurysms. Scale 50 µm.

### All neurites in the senile plaques display some degree of Tau phosphorylation

Tau phosphorylation is another important pathological hallmark of Alzheimer’s disease. Previous studies showed that Tau phosphorylation is tightly linked to Aβ-related lysosome destabilization[42, 44]. If Aß deposition induces lysosome destabilization in most neurites in senile plaques, we should expect to see the whole senile plaque marked with Tau phosphorylation. However, in previous studies, only some neurites in senile plaques showed phos-Tau-labeled neuritic dystrophy. Did we miss some weak phos-Tau signals in senile plaques in previous studies? We re-examined Phos-Tau staining in senile plaques with longer exposure. We noticed that there existed two type of phos-Tau staining in senile plaques (Figure 7). A strong type of phos-Tau staining resembled dystrophic neurites as described before. Another weaker type of phos-Tau staining essentially labeled all the rest of neurites in the senile plaques. Co-localization analysis with ImageJ showed that 34±5% (N=6) of phos-Tau staining colocalized with senile plaque MetaBlue blue autofluorescence (Example images of colocalization analysis was provided in Supplementary Figure 2). Tau phosphorylation could be a dynamic event and be quickly attenuated in some *in vivo* conditions and the weak type of Tau phosphorylation probably goes unnoticed in many cases.

**Figure 7.**
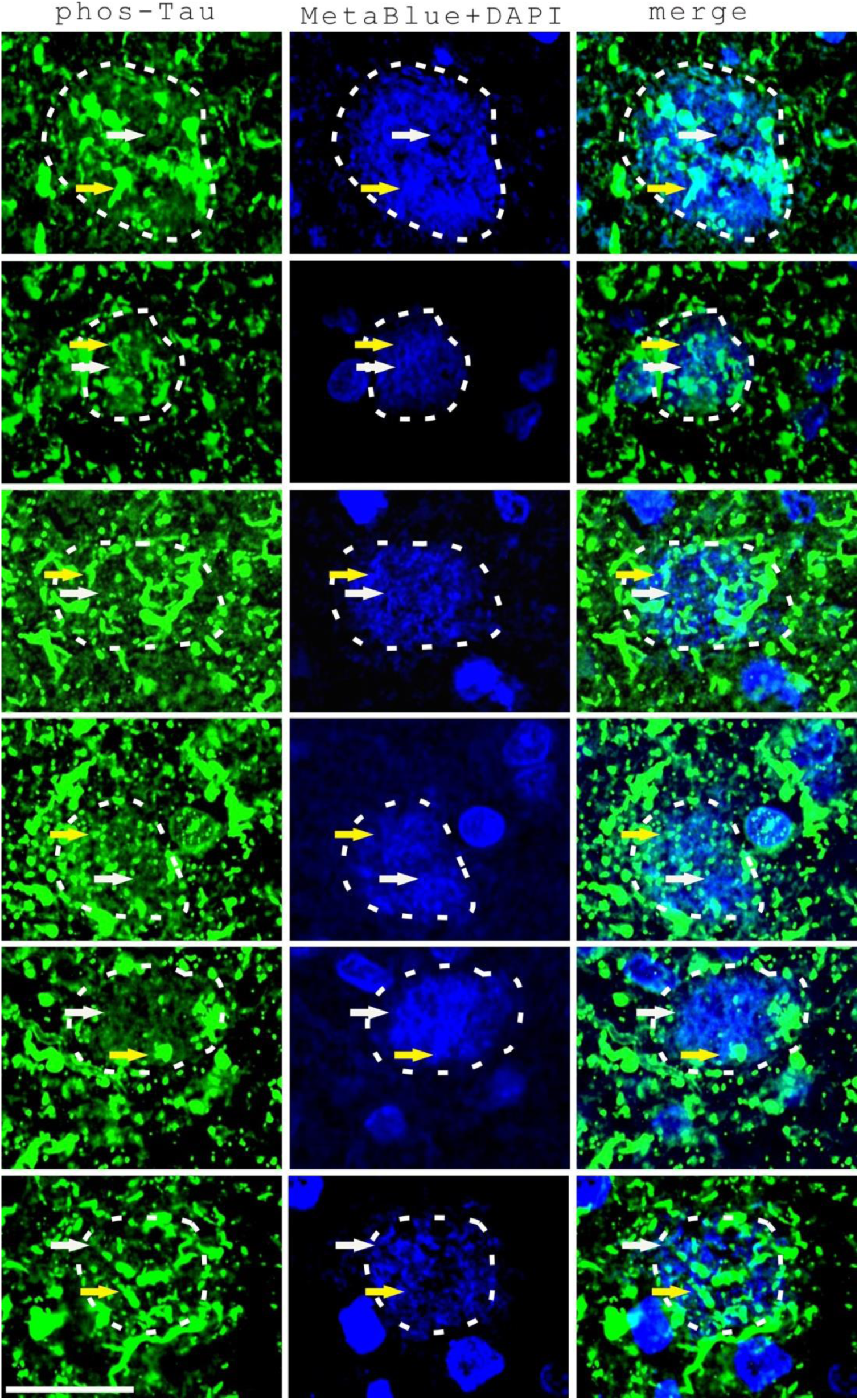
Almost all neurites in senile plaques in AD frontal brain tissues were labeled by phos-Tau immunostaining either with strong signals or with weak positivity. The dashed lines indicate the senile plaques marked with amyloid-related blue autofluorescence. The yellow arrows indicate dystrophic neurites with strong phos-Tau staining (with green fluorescence) while the white arrows indicate neurites with weak phos-Tau staining. Co-localization analysis with ImageJ showed that 34±5% (N=6) of phos-Tau staining colocalized with MetaBlue blue autofluorescence in the senile plaques. Scale, 25 µm.

## Discussion

Senile plaque is the most prominent pathological hallmark in AD. The pathological mechanism of senile plaque formation in AD remains unsolved for over than one hundred years. It is an awkward situation when we are explaining AD therapies to patients without a clear pathological mechanism of senile plaque pathogenesis. The research on AD in general has become increasingly fragmented with over 260,000 papers with the keyword of “Alzheimer’s disease” in the PubMed database, which makes it exceedingly difficult to put the “jigsaw puzzle” of AD into one piece. However, we believe that, it is very important to make effort to build some theoretic common grounds in this prolonged battle against AD.

This study attempts to provide a simple, reasonable explanation to the mechanism of senile plaque formation in AD. The model can be summarized as the following: senile plaques are mainly formed by destabilized-lysosome-enriched neural processes catabolizing hemorrhagic amyloid materials. The evidence is summarized below: Firstly, senile plaques are mainly consisted of degenerative dystrophic neurites with Aβ deposition and significantly-reduced MAP2 staining, which links senile plaque formation with neurite degeneration directly. It appears that Aβ deposition induces beading-type MAP2 degeneration in neurites in senile plaques. This result is consistent with previous studies indicating that Aß induces beading-type neurite degeneration in mouse N2a or rat hippocampal neuron culture *in vitro* experiments[45, 46]. Secondly, senile plaques can be labeled by lysosome marker Cathepsin D (Figure 2), endo-lysosome marker Sortilin 1 (Figure 3) and also Lamp2 (Supplementary Figure 3) with dual-component granule type and diffusive type staining patterns. The diffusive type endo-lysosomal marker staining indicates endo-lysosome destabilization as suggested from previous studies [42, 44]. Quantitation showed that around 85% of Aβ was in the areas with diffusive Cathepsin D staining, highlighting that destabilized lysosomes are the major locations of neuritic amyloidosis, which is consistent with previous studies indicating that Aβ is a lysosome destabilizer[22, 47]. Lysosome marker Lamp1 staining in senile plaques was previously studied in mouse models extensively [21, 48, 49]. The enrichment of Lysosome-related markers such as Lamp1, Cathepsin D, PLD3 and Sortilin 1 in human brain tissues have also been reported [50–53]. In this study, the dual-component staining patterns of Aβ, Cathepsin D, Sortilin 1 in relation to lysosome destabilization in senile plaques were characterized in detail, which has not been done in previous studies, to our best knowledge. Thirdly, senile plaques also colocalized with hemorrhagic markers such as Ceruloplasmin, Hemin, HBA, HbA1C, ApoE (Figure 4 and Supplementary Figure 3). The association of Ceruloplasmin and ApoE with senile plaques has been studied previously[54, 55]. We noticed that all these hemorrhagic-related markers colocalized with Aβ and showed similar dual-component granule type and diffusive type staining as Cathepsin D and Sortilin 1, suggesting they are also existing in endo-lysosomal or destabilized endo-lysosomal compartments in senile plaques. In summary, Aβ deposition in diffusive senile plaques is mainly happening intracellularly in degenerative neuronal processes containing mostly destabilized endo-lysosomal vesicles and hemorrhagic amyloid materials.

This model suggests that senile plaques are simply clustered neural processes internalizing hemorrhagic amyloid materials such as Aβ, HBA etc. We did an extra analysis by measuring mean Aβ intensities in individual Aβ-positive axons and senile plaques. The average Aβ intensity of individual Aβ-positive axons (N=120) showed very similar mean Aβ staining intensity (97±13%, p=0.459, Mann Whitney test) as the mean intensity of Aβ in senile plaques (N=18). A picture comparing Aβ deposition in individual axons, in small axon clusters and in senile plaque axons was shown in Supplementary Figure 4. Based on our estimate, by dividing mean area intensity of senile plaque Aβ on the average area intensity of axonal Aβ, a senile plaque might contain as many as 60.62±39.18 Aβ-positive axons. The actual axon number could be lower since senile plaques contain not only axons but also dendritic processes. Thus, senile plaques indeed can be considered as highly-clustered Aβ-positive neural processes.

Why Aβ-positive neural processes are so concentrated in senile plaques? There could be two possible explanations: Firstly, as the brain tissue is heavily enriched with neural processes such as axons and dendrites, when senile plaque formation is triggered by microaneurysm rupture, a large amount of Aβ released from the rupture will strongly affecting Aβ-deposition in multiple neighboring neural processes simultaneously. Secondly, vascular/hemorrhagic amyloid spill might sometimes hit axon bundles containing many axons in clusters, which is a special anatomic feature of axon arrangement in the brain[42]. The morphogenesis of senile plaque is likely affected by the position, strength and content of hemorrhagic leakage, vascular degeneration, the distribution of axons and dendrites and also their interactions with glial cells.

Tau phosphorylation in neurofibrillary tangles and neuropil threads is another pathological hallmark of AD. Our previous study indicated that lysosome destabilization mediates the induction effect of Aβ deposition on Tau phosphorylation[42]. If senile plaques are mostly consisted of Aβ-deposited neural processes with lysosome destabilization, we should expect to observe a wide-spread Tau phosphorylation in senile plaques instead of just a few dystrophic neurites. Indeed, when Tau phosphorylation staining in senile plaques is imaged with longer exposure settings, most neurites in the senile plaques are positive with Tau phosphorylation staining (Figure 7). However, some of the Tau phosphorylation staining on degenerative neurites in senile plaques is very weak. Similarly, strong or diffusive, fragmented and weak Tau phosphorylation signals could also be observed in neurons or neuropil threads (Supplementary Figure 5). Many of these weak pho-Tau signals could be ignored under conventional staining and imaging conditions. The significance of these weak phos-Tau signals and the dynamic change of Tau phosphorylation intensity is worthy of further investigation.

This newly proposed model can be aligned with previous theories and findings in a logical way. The theory of senile plaque formation by hemorrhagic material-enriched destabilized lysosomes is not in conflict but rather enriches the contents and further advances most previous theories, such as the amyloid cascade hypothesis, vascular degeneration, mitochondria failure, neural inflammation etc. The importance of Aβ in AD neurodegeneration stated in the amyloid cascade hypothesis[6] is supported with the presented data. However, the original assumption that Aβ comes from a neuronal source is probably incorrect because mounting evidence indicates that Aβ peptides in senile plaques might mainly come from hemorrhagic amyloid material leakage into the brain parenchyma, colocalizing with multiple blood or vascular, glial foot markers such as Hemin, HbA1C, Hemoglobin, ApoE, LRP1, ColIV, GFAP[14]. However, these amyloid peptides are largely processed in the neurons intracellularly. Early studies suggests that amyloidogenic Aβ production might happen in lysosomes[6, 56]. This study emphasized that Aβ amyloidosis does happen in lysosomes, however, with an important modification, mostly happen in destabilized lysosomes. In addition, the importance of Aβ is probably underestimated in previous models. Aβ not only affects neurodegeneration but also likely impact on earlier pathological stages in AD pathogenesis, such as intravascular hemolysis and vascular degeneration, in which lysosome destabilization could also be involved. Aβ internalization and lysosome destabilization is predicted to induce proteomic damage to the neural processes, such as axon, dendrite or synapse damage, so that synaptic toxicity [15, 16] is a natural result of neural cell amyloidosis. Mitochondria deficiency can be triggered by both oxidative stress resulting from hemolysis, vascular degeneration or mitochondria damage due to intracellular amyloidosis-related proteomic injury[17, 18]. Vascular degeneration, blood leakage and neurite damage also certainly induces neural inflammation [23, 24]. Metal ion toxicity [25–27] can be easily explained by the abundant metal-containing plasma proteins in the blood leakage. For example, hemoglobin is an iron-containing protein and Ceruloplasmin is copper-binding protein. Albumin and transferrin are known as major binding proteins for aluminum so that aluminum enrichment in senile plaques could also be explained. Metal ions such as iron and copper can also induce oxidative stress with the production of lipid peroxidation products such as MDA and 4HNE, which could further induce DNA damage. Oxidative stress could also be resulted from mitochondria deficiency and further amplifies mitochondria damage. Since lysosome is the end organelle for autophagy and endosome pathway, it is not surprising to find other autophagy or endosome related proteins trapped in senile plaques in the form of destabilized lysosome components. Based on this model, we predict that many materials involved in hemolysis, cerebral vascular degeneration, immune reactions to hemolysis and cerebral vascular degeneration, autophagy, endocytosis and the lysosomal pathway could end-up in the senile plaques and be capable of labeling senile plaques efficiently. Recent publications indicated that a group of cerebrovascular extracellular matrix related proteins called matrisome proteins could be novel markers of senile plaques[57, 58]. Our suggested model can explain why these cerebrovascular proteins accumulate in senile plaques. We believed that amyloid accumulation and lysosome destabilization could be a way of transcellular stress inheritance so that inflammation and tissue damage can be passed on from one cell type to another cell type through endocytosis and lysosome destabilization. Regarding the central role of lysosome destabilization in Alzheimer’s disease, AD could be considered as a special type of lysosome storage disease, a lysosome storage disease induced by the overburdened neuronal lysosome processing of chronic hemorrhagic and vascular amyloid material leakage. A previous study stated that senile plaques are formed by individual neurons experiencing faulty autolysosome acidification-induced autophagic Aβ build-up and cell death in Alzheimer’s disease mouse models[20]. We agree that there is serious lysosome defects in AD neurons, however, the existing of a central degenerating nuclei resulting from neuronal cell death in the cores of senile plaques can not be confirmed with our experiments. Instead, we proved that all the cores of dense-core senile plaque don’t have nuclei but instead possess unique blue autofluorescence derived from Aß self-oligomers or hetero-oligomers[41].

We want to emphasize that there are some significant difference in this model vs. previous theories. Firstly, we think that senile plaque Aß comes from blood/vascular Aß leakage instead of totally deriving from neurons themselves. Although there are many excellent studies indicating the potential link of vascular/blood system with AD[10–13, 32, 59, 60], the vascular/ blood origin of senile plaques as a theory is not firmly established. Our previous and current studies gathered critical evidence that both dense-core senile plaques and diffusive senile plaques are tightly associated with hemorrhagic markers such as HBA, Hemin, HbA1C and Ceruloplasmin. On the other hand, the logic of neuron-produced Aß inducing Alzheimer’s disease is not flawless. It is very hard to understand that neurons, the “smartest” type of cells in the body, will intoxicate themselves by continuously self-producing toxic Aß, completely lacking a biological negative feedback mechanism. It is also puzzling when CCA with abundant Aß is frequently observed in AD but the final whereabouts of CCA Aß is seldom discussed or explained. Our model stating blood leakage of Aß intoxicates neurons through a chronic “lysosome storage” mechanism can explain the pathological findings well. In this model, CAA Aß will leak into the brain mesenchyme through microaneurysm rupture to induce senile plaque formation. Secondly, our model highlights the crucial importance of axonal defects in AD while many AD researches think the problem in AD is in the defects of neuron cell bodies. This new model suggests that neuritic damage is a direct result of senile plaque Aß deposition as the “first hit” in AD, although Aß deposition in neurites might have a damaging effect on the neuronal cell bodies, providing the “second hit”, in the long run.

This new model provokes rethinking of the current treatment and prevention strategy for AD. For disease modification therapy of AD, limited success with lecanemab[36] and donanemab[37] is achieved. Disease modification therapies targeting Tau is also under development[61–64]. Our data suggest that both Aß and Tau are linked to intracellular events such as lysosome destabilization. Apparently, both Aß and Tau affects not just individual neurons but instead affects the brain at the network level through numerous neuronal connections with axons and dendrites (Supplementary Figure 6 and 7). To reduce extracellular amyloid is predicted to only partially help on the recognition recovery of AD patients since the neuron function is severely negatively affected at a network level by the intracellular amyloid deposition. A recent research indicated that there was a potential discrepancy between Aß PET signal reduction and histo-pathology improvement and the deep layer brain pathology hardly changed during the aducanumab antibody treatment in AD patients[65].Similarly, the obstacle of Tau targeting approach is how to deliver the Tau inhibitors into the neural cells efficiently. Both amyloid and Tau therapies will be affected by the timing of the therapy and the route of delivery of the therapeutic reagents. Another important target of AD therapy might be the lysosomes. This study indicated AD has similar “lysosome storage” characteristics of other classical lysosome storage diseases. Thus, restoring lysosomal functions can be beneficial to both AD and lysosome storage disease. On the other hand, the data suggested that a promising approach of AD prevention is by reducing vascular and blood pathologies since senile plaques are induced by hemorrhagic events. If we can successfully prevent chronic hemolysis, vascular degeneration and microhemorrhage, probably, AD can be largely prevented.

Lysosome destabilization is a crucial event in AD pathogenesis, which mediates the effect of Aβ on Tau phosphorylation and has a direct effect on neuronal degeneration. The destabilization of lysosomes not only potentially leaks lysosome enzymes but also lysosome enzyme substrates and degradative intracellular or extracellular-originated peptides or lipid components or other biological macromolecules into the cytoplasm, which has the potential to unpredictably change the behaviors of hundreds or even thousands of molecules both from intracellular source and extracellular source and countless unforeseeable chain reactions. The scenario is further complicated with the existence of several important lysosome repair pathways[66–68], which means, under certain conditions, lysosome damage is reversible. How chronic lysosome leakage induces neural cell damage and dementia in AD will become a critical topic of future investigation.

## Acknowledgements

We want to thank the help from Dr. Ma Chao and Dr. Qiu Wenying for providing AD tissue sections from National Human Brain Bank for Development and Function, Chinese Academy of Medical Sciences and Peking Union Medical College, Beijing, China. Additionally, we want to thank Prof. Ming Yang from Shanghai Jiao Tong University, Dr. Peng Du and Dr. Lei Xia from Xinhua Hospital for valuable scientific discussions. Moreover, we want to thank Housheng Wang, Pingxin Liu, Xiaoyi Bao, and Jun Wang from Shanghai Jiao Tong University for excellent laboratory assistance.

## Supplementary Figures

**Supplementary Figure 1.**
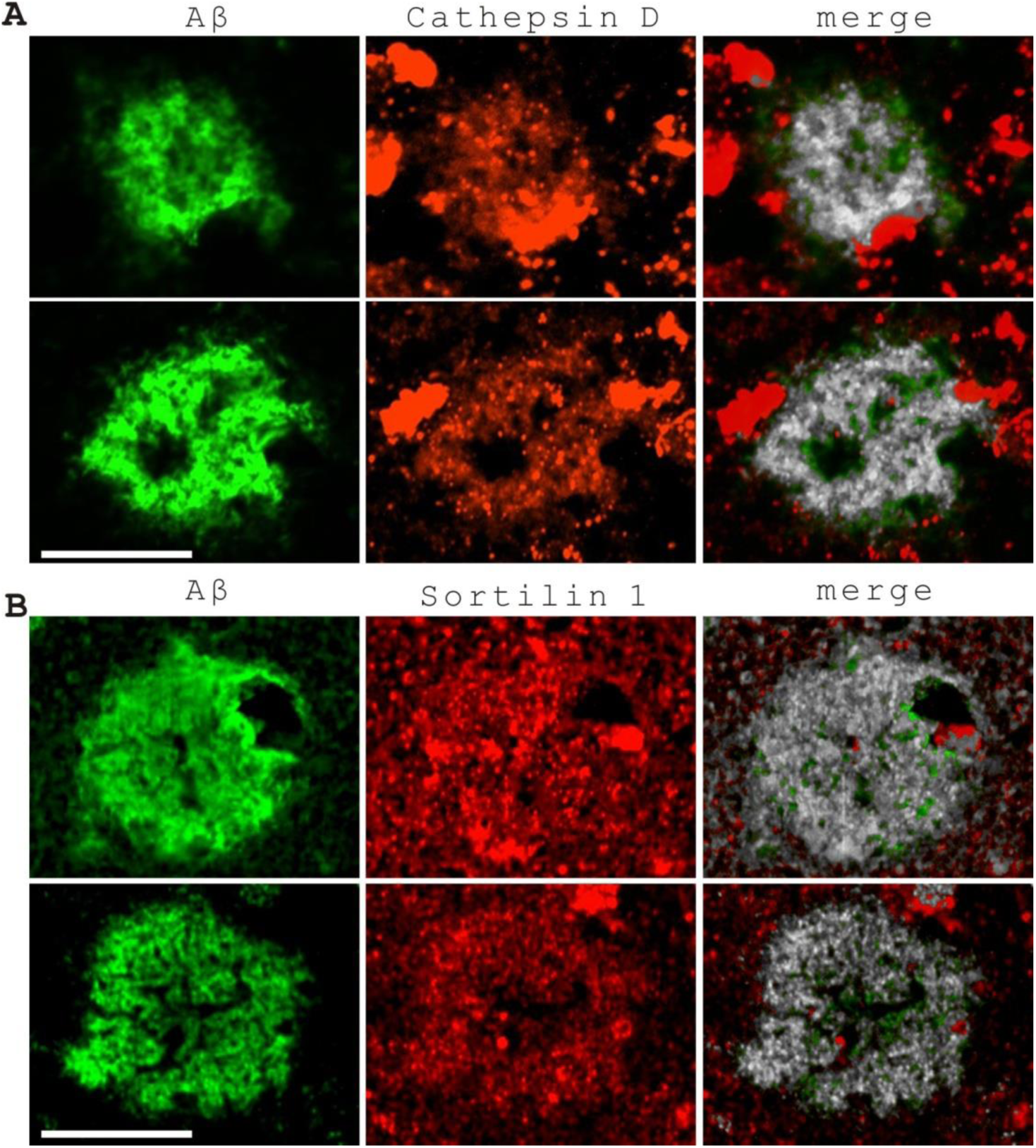
Colocalization analysis of Aβ and lysosome-related markers in senile plaques in AD frontal brain tissues. Colocalization analysis was performed using the ImageJ “Colocalization Threshold” plugin. AD: Alzheimer’s disease. (A) Colocalization analysis of Aβ and Cathepsin D in senile plaques. Colocalization between Aβ (green) and Cathepsin D (red) is highlighted with white pixels. The analysis estimated that 74% (top panel) and 84% (bottom panel) of Aβ staining colocalized with Cathepsin D staining in these two samples, respectively. (B) Colocalization analysis of Aβ and Sortilin 1 in senile plaques. Colocalization between Aβ (green, Alexa Fluor 488) and Sortilin 1 (red, Alexa Fluor 594) is highlighted with white pixels. The analysis estimated that 88% (top panel) and 83% (bottom panel) of Aβ staining colocalized with Sortilin 1 staining in these two samples respectively. Scale bar: 25 μm.

**Supplementary Figure 2.**
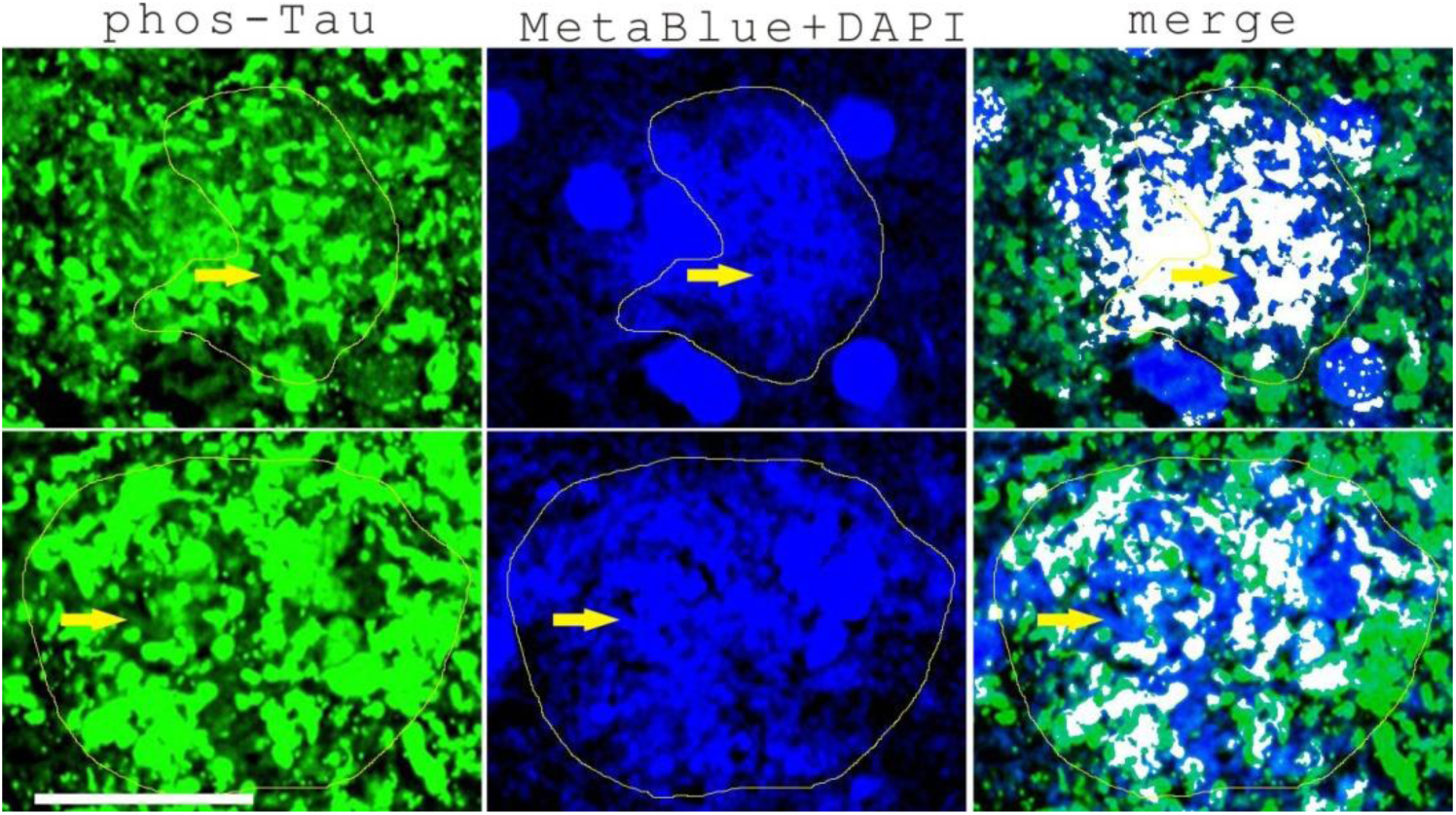
Two set of images showing the colocalization analysis of Tau phosphorylation and senile plaques (labeled with MetaBlue, amyloid blue autofluorescence) with ImageJ colocalization threshold plug-in. The analysis indicated 58% (top) and 45% (bottom) of phos-Tau colocalized with senile plaque blue autofluorescence. The colocalization areas were highlighted with white pixels. The yellow lines indicated the senile plaque areas. The yellow arrows indicated that the weak phos-Tau signals were often not picked up by the software analysis so that the actual colocalization ratio between phos-Tau with senile plaque blue autofluorescence was underestimated. Scale bar, 25 µm.

**Supplementary Figure 3.**
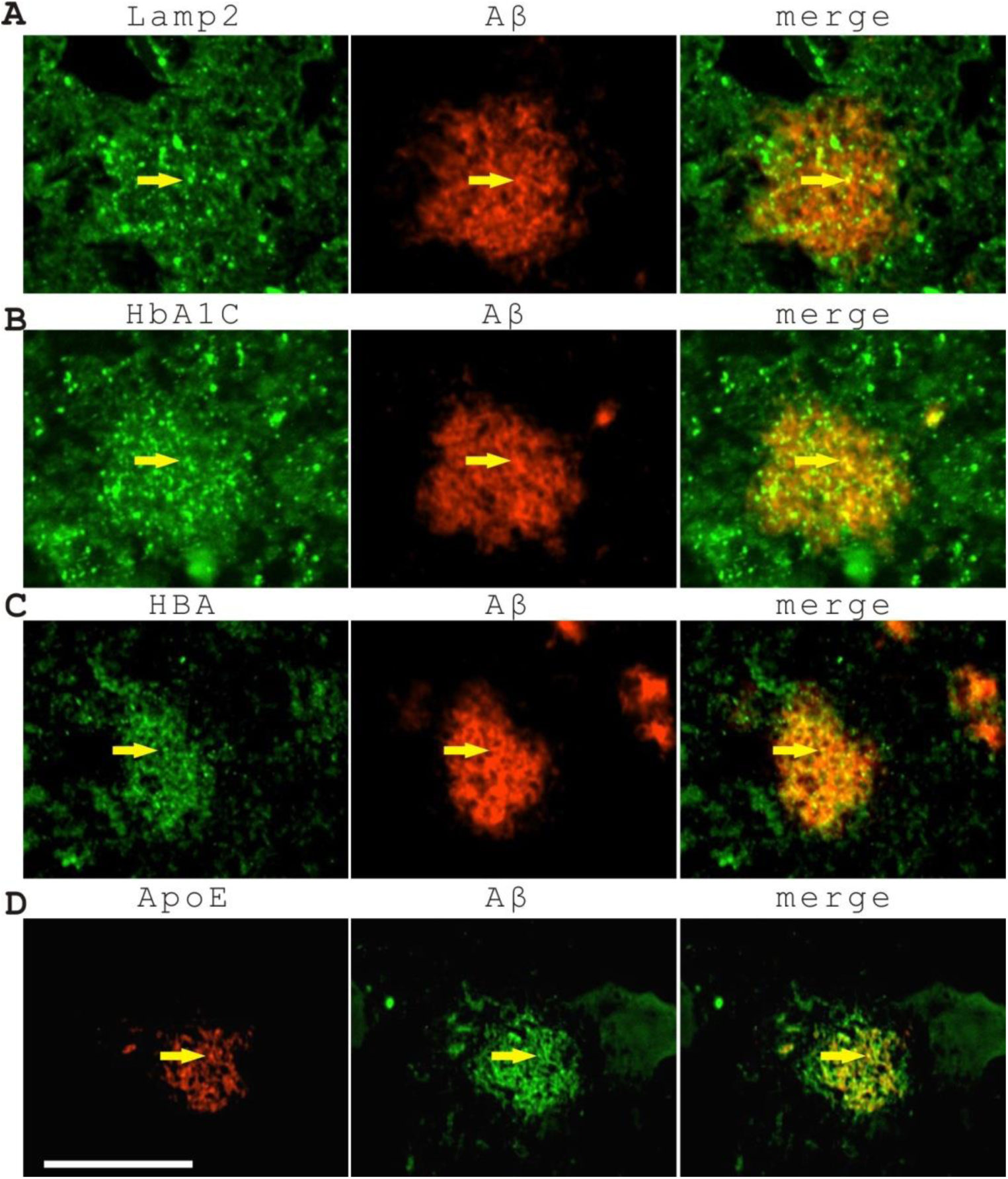
Lamp2, HbA1C, HBA, ApoE staining of AD front brain tissues all showed “sesame cookie” type of staining with variable number of granules and also diffusive neuritic staining in the senile plaques. (A) Lamp2. (B) HbA1C. (C) HBA. (D) ApoE. The arrows indicated the granule type marker staining. Scale bar, 25 µm.

**Supplementary Figure 4.**
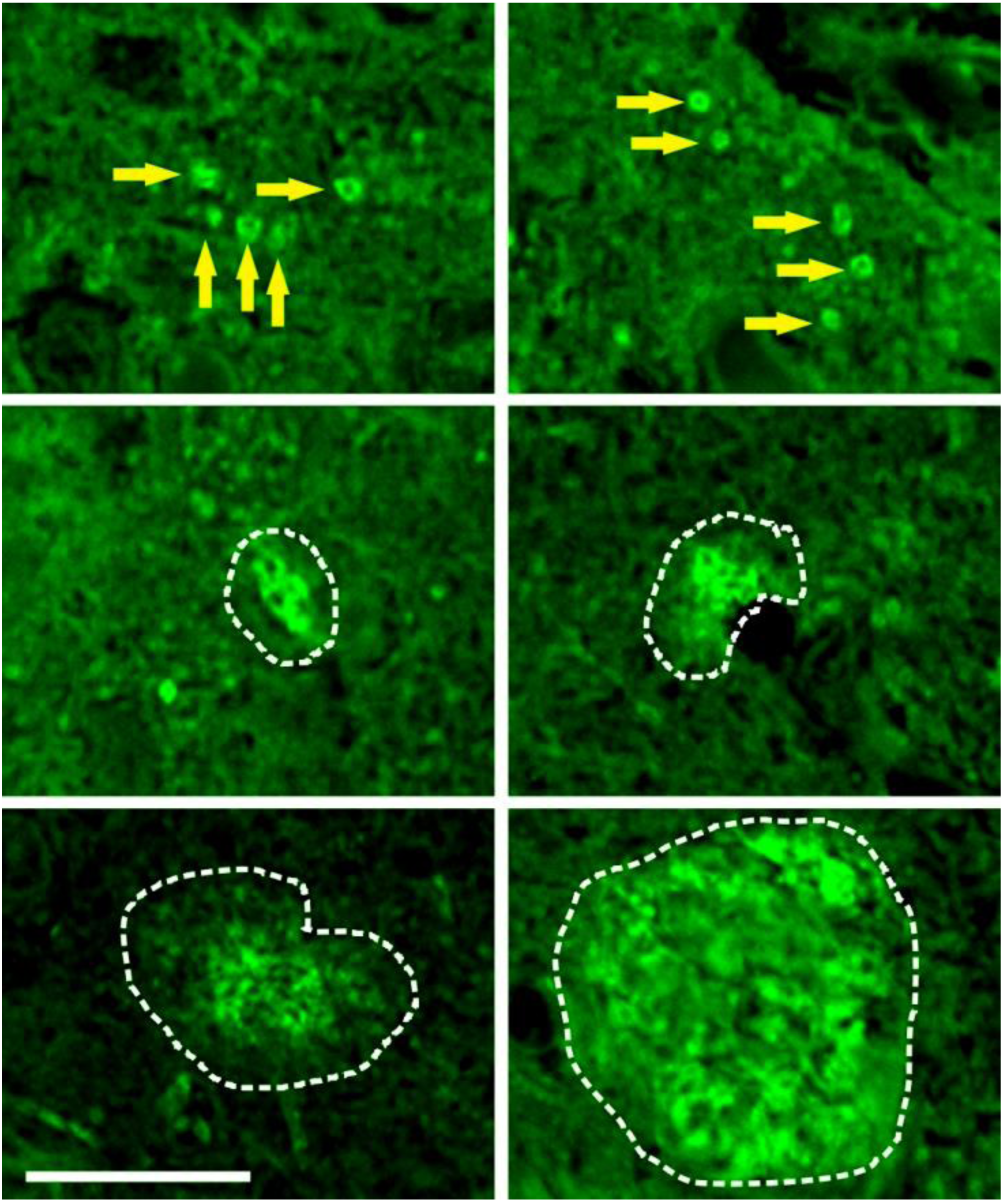
Senile plaques are clusters of Aβ-positive neural processes (mostly axons) as shown by Aβ antibody staining of AD frontal brain tissues. Top panel showed Aβ-labeled scattered individual axons. The middle panel showed two very small clusters of Aβ-positive axons. The bottom panel showed larger senile plaques containing many Aβ-positive axons. The average Aβ intensity of Aβ-positive axons (N=120) showed very similar mean Aβ staining intensity (97±13%, p=0.459, Mann Whitney test) as the mean intensity of Aβ in senile plaques (N=18). Scale bar, 25 μm.

**Supplementary Figure 5.**
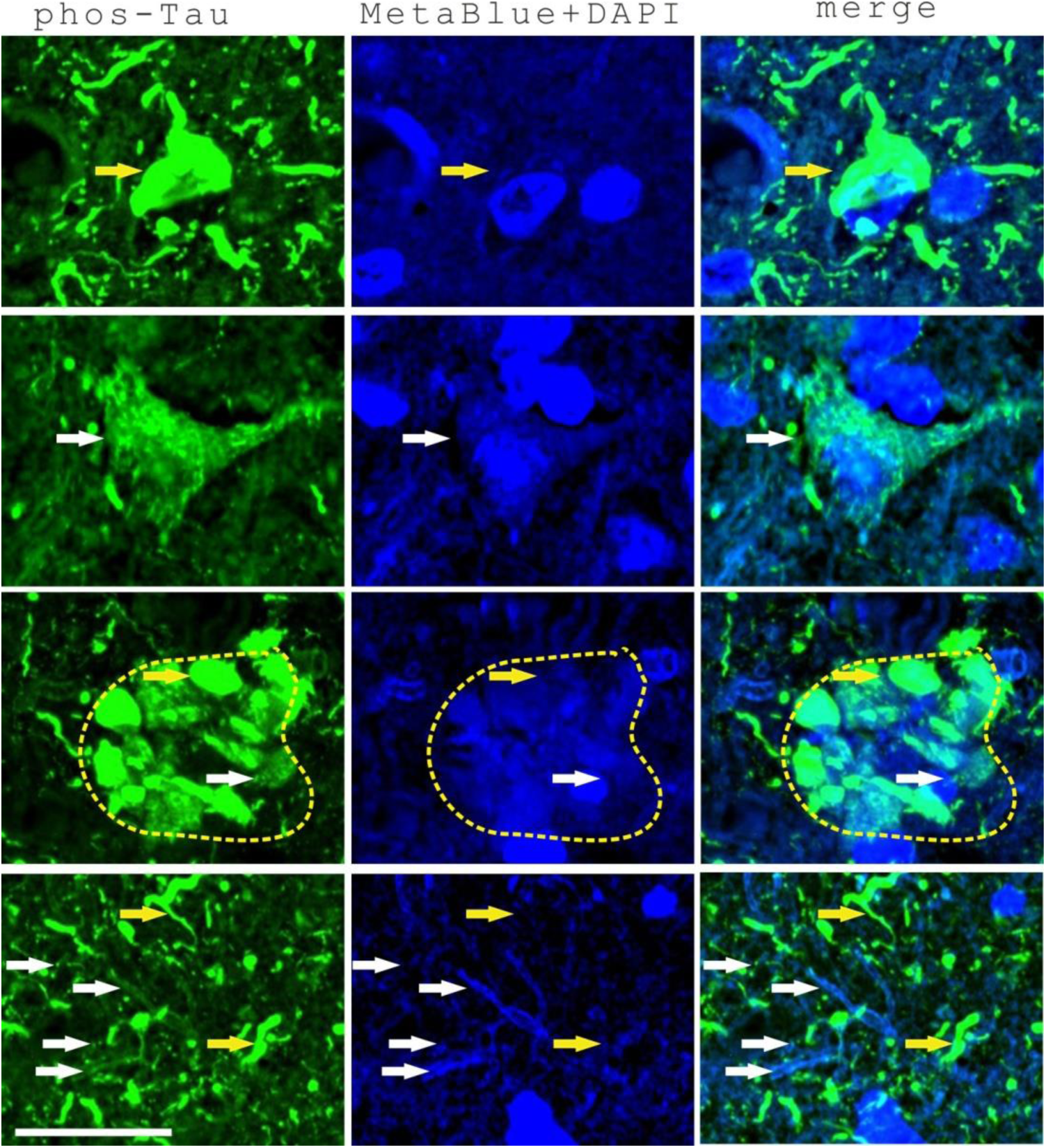
Both strong and weak staining patterns of Tau phosphorylation have been observed in the neural cells, senile plaques and neurites in AD frontal brain tissues, as shown by phos-Tau antibody immunostaining. The top row indicated a tangled neuron with strong, typical phos-Tau staining. The second row indicated a neuron with non-typical weak, fragmented and diffusive phos-Tau staining in the cell soma. The third row showed strong (yellow arrow) or weak dystrophic staining (white arrow) of phos-Tau in the senile plaque region. The bottom row showed both strong neuritic phos-Tau staining (yellow arrows) and also weak and diffusive or fragmented neuritic phos-Tau staining (white arrows). Scale bar, 25 µm.

**Supplementary Figure 6.**
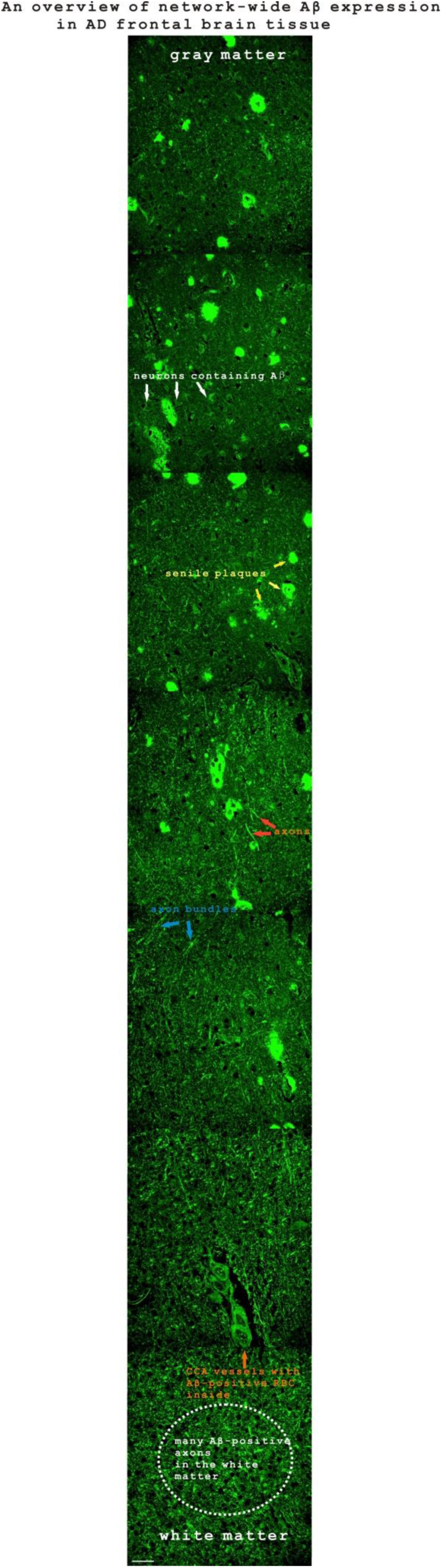
An overview of network-wide expression of Aβ, shown by Aβ immunostaining in an AD frontal brain tissue sample. The white arrows indicated several neurons with Aβ staining. The yellow arrows indicated several senile plaques. The red arrows pointed to two individual axons with Aβ staining. The blue arrows indicated two Aβ-positive axon bundles. The orange arrow indicated a blood vessel with Aβ staining. The white dashed circle indicated many Aβ-positive axons in the white matter. Scale bar, 50 µm.

**Supplementary Figure 7.**
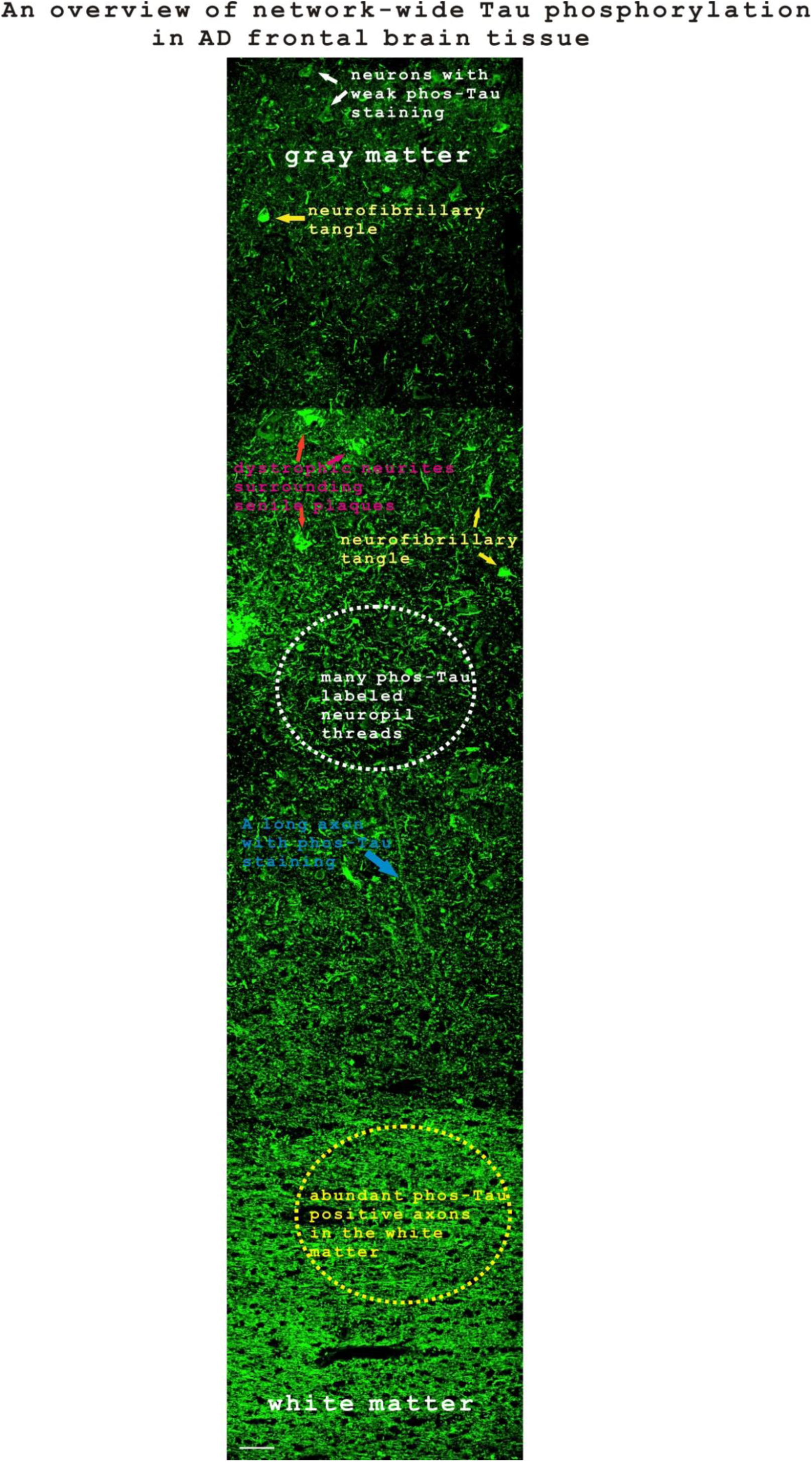
An overview of network-wide expression of Tau phosphorylation in an AD frontal brain tissue sample as shown by phos-Tau immunostaining. The white arrows indicate two neurons with weak phos-Tau staining. The yellow arrows indicated several neurofibrillary tangles. The red arrows indicated three clusters of dystrophic neurites surrounding senile plaques. The white dashed circle indicated abundant neuropil threads. The blue arrow indicated a long axon positive with phos-Tau staining. The yellow dashed circle indicated many phos-Tau-positive axons in the white matter. Scale bar, 50 µm.

## Notes

### Competing Interest Statement

The authors have declared no competing interest.

